# The brain’s sensitivity to sensory error can be modulated by altering perceived variability

**DOI:** 10.1101/2023.06.26.546615

**Authors:** Ding-lan Tang, Benjamin Parrell, Sara D. Beach, Caroline A. Niziolek

## Abstract

When individuals make a movement that produces an unexpected outcome, they learn from the resulting error. This process, essential in both acquiring new motor skills and adapting to changing environments, critically relies on error sensitivity, which governs how much behavioral change results from a given error. Although behavioral and computational evidence suggests error sensitivity can change in response to task demands, neural evidence regarding the flexibility of error sensitivity in the human brain is lacking. Critically, the sensitivity of the nervous system to auditory errors during speech production, a complex and well-practiced motor behavior, has been extensively studied by examining the prediction-driven suppression of auditory cortical activity. Here, we tested whether the nervous system’s sensitivity to errors, as measured by this suppression, can be modulated by altering speakers’ perceived variability. Our results showed that error sensitivity was increased after exposure to an auditory perturbation that increased participants’ perceived variability, consistent with predictions generated from previous behavioral data and state-space modeling. Conversely, we observed no significant changes in error sensitivity when perceived variability was unaltered or artificially reduced. The current study establishes the validity of behaviorally modulating the nervous system’s sensitivity to errors. As sensitivity to sensory errors plays a critical role in sensorimotor adaptation, modifying error sensitivity has the potential to enhance motor learning and rehabilitation in speech and, potentially, more broadly across motor domains.

**Significance Statement:** The process of learning from error is essential for both the acquisition of new skills and successful adaptation to changing environments. Such error-based learning critically relies on error sensitivity, which determines how much we learn from a given error. Although evidence from behavioral studies suggests error sensitivity is malleable, neural evidence regarding the flexibility of error sensitivity in the human brain is lacking. Here, we showed that the nervous system’s sensitivity to errors can be modulated by altering perceived variability. The present study establishes the validity of behaviorally modulating neural sensitivity to sensory errors. Improving our ability to learn from error can play a critical role in applied settings such as rehabilitation.

## Introduction

Throughout our daily activities, we constantly monitor the outcomes of our actions and adjust our behavior when these outcomes don’t match our expectations. For example, when throwing a ball, the outcome will be affected by changes in both external conditions (e.g., the strength and direction of the breeze) and internal conditions (e.g., muscle fatigue). If these conditions cause the ball to end up in a different place than we expect, we unconsciously adjust our next throw to correct for this error. A critical variable in such *sensorimotor learning* is error sensitivity, which is the “learning rate” determining how much is learned from trial to trial (Thoroughman & Shadmehr, 2000). When perturbations are consistent (like a steady breeze when throwing the ball), correcting for errors allows us to *adapt* to these new conditions. The same error-correction system also operates in the absence of external perturbations, both correcting for and contributing to variability in our actions across repetitions (Ahn et al., 2016; Blustein et al., 2021; van Beers, 2012).

Early models of error-based sensorimotor adaptation assumed that the sensitivity to error for a given action is constant; that is, the brain always learns a certain fraction of the error (Cheng & Sabes, 2006; Scheidt et al., 2001; Smith et al., 2006; Thoroughman & Shadmehr, 2000; van Beers, 2012). However, recent work using behavioral psychophysics and computational modeling of reaching movements has suggested that error sensitivity is not static, but changes as a function of error size (Marko et al., 2012) and the history of previously experienced error (Herzfeld et al., 2014). Similarly, although movement variability is often thought to be constant, arising largely from unwanted noise in the nervous system (Faisal et al., 2008), recent work has demonstrated that motor variability is more actively controlled (Tang et al., 2022; Wong et al., 2009). Importantly, changes in variability in reaching also affect adaptation (Wu et al., 2014), consistent with error sensitivity playing a shared role in both processes. Despite this evidence from behavioral analyses that error sensitivity in the motor system is flexible, there is a paucity of neural evidence regarding whether and how error sensitivity can be modulated in the human brain.

Speech production provides a unique opportunity to examine changes in neural sensitivity to errors during motor behavior. The sensitivity of the nervous system to auditory errors in speech has been extensively studied by examining the suppression of the auditory cortical response to self-produced speech compared with its response to playback of the same speech signal (e.g. Curio et al., 2000; Flinker et al., 2010; Houde et al., 2002; Ventura et al., 2009). This speaking-induced suppression (SIS), primarily seen in the left hemisphere, is thought to reflect a partial neural cancellation of incoming auditory feedback by efference copy prediction. Critical for our purposes, SIS has been shown to be a neural marker of sensitivity to sensory error (Behroozmand & Larson, 2011; Chang et al., 2013; Niziolek et al., 2013; Sitek et al., 2013): that is, the larger the prediction error perceived, the smaller the suppression.

In the current study, we use SIS to test whether the nervous system’s sensitivity to errors can be modulated by auditory feedback perturbations that alter speakers’ perceived variability. In a previous behavioral study (Tang et al. 2022), participants adjusted their produced variability in response to both an inward-pushing perturbation that reduced their perceived trial-to-trial variability (Figure 1A, left; all productions shifted towards the center) and an outward-pushing perturbation that increased their perceived trial-to-trial variability (Figure 1A, right; all productions shifted away from the center). Critically, simulations of these perturbations with a state-space model of learning suggested that the behavioral response to the outward perturbation was likely caused by an increase in speakers’ sensitivity to auditory error, which drove trial-to-trial overcorrections that increased variability. Based on these results, we predicted that exposure to the outward perturbation would result in an increase in error sensitivity that would be reflected in a reduction in SIS in this condition. Such suppression in the auditory domain provides a window into sensory error processing during human motor control. As sensory error is the driver of sensorimotor adaptation, understanding how to modulate the sensitivity to that error in the human brain enables us to probe the fundamental mechanisms of sensorimotor control. Moreover, it has the potential to enhance motor learning and rehabilitation in speech and, potentially, other motor domains.

**Figure 1.**
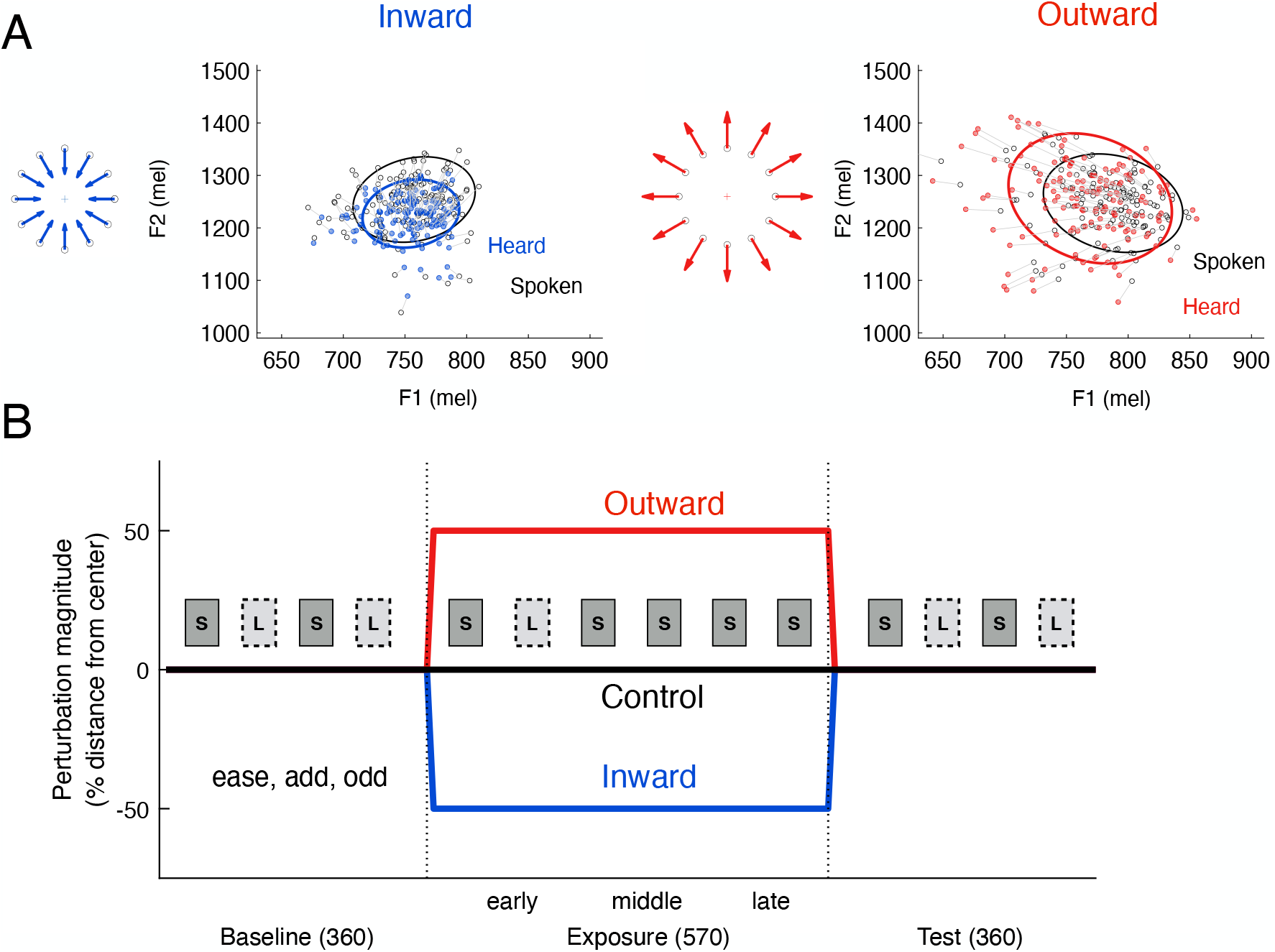
Experiment design. **A**. Schematic and examples of perturbations (inward and outward) applied to vowel formants during the exposure phase from a single representative participant. The formant values that the participant produced and heard are indicated by black and colored circles, respectively. The ellipses represent a 95% confidence interval around the data points of the same color. All auditory formant perturbations were applied in mels, a logarithmic measure of frequency. **B**. Experimental procedure. Each participant completed three sessions, one for each of the three perturbations (inward, outward, control) applied during the exposure phase. Participants performed both a speaking task (“S”) and a listening task (“L”), grouped into blocks of 90 trials. The stimulus words were “ease”, “add” and “odd”, pseudo-randomized within each block.

## Results

### Speech variability can be modulated by alterations of perceived acoustic variability

Previous work in speech production suggests that an outward auditory feedback perturbation, which increases perceived variability, results in increases in produced variability due to increased sensitivity to auditory errors (Tang et al. 2022). We conducted an initial behavioral study (pilot group: 10 participants) to confirm that the addition of passive listening trials, necessary for our neural metric of auditory error sensitivity, does not alter the behavioral response to this outward perturbation (Figure 1B, outward condition only). In speaking blocks, participants produced the stimulus words (“ease”, “add” and “odd”) one at a time while receiving auditory feedback of their own speech through circumaural headphones. Blocks of listening trials were interspersed between speaking blocks, as would be required for measuring SIS. Vowel formants, resonances of the vocal tract that distinguish between different vowels, were perturbed during the exposure phase, with formants pushed outward from the center of each vowel (50% of the distance to the center in 2D mel frequency space). Vowel formants were unperturbed in initial baseline and final test phases. As in our previous study, participants significantly increased their produced variability during the outward perturbation (Figure S1; main effect of phase: F(2.1,18.7) = 4.741, p = 0.021, η_p_^2^= 0.345; +6.1 mels in late exposure phase, t = 2.86, p = 0.05, d = 0.91), verifying the feasibility of the current experimental design to induce behavioral changes in produced variability.

For the main MEG study, each participant (N=15) took part in three separate MEG sessions, each separated by at least one week, during which they received different auditory feedback during the exposure phase of the experiment (Figure 1B): an inward perturbation that decreased perceived variability, an outward perturbation that increased perceived variability, and a no-perturbation control (normal auditory feedback). There was no significant difference in baseline variability across the three sessions (F(1.2,17.6) = 0.872, p = 0.388), suggesting movement variability during speech production remains stable over time in the absence of any auditory perturbation.

We then measured how participants changed their produced variability in response to the feedback perturbations. Produced variability was stable in the control session (Figure 2, no main effect of phase: F(2,28) = 1.006, p = 0.378), while participants significantly increased their produced variability during both the inward (Figure 2, main effect of phase: F(2,28) = 4.545, p = 0.02, η_p_^2^= 0.245) and outward (main effect of phase: F(1.4,19.9) = 4.829, p = 0.029, η_p_^2^= 0.256) perturbation sessions. Post-hoc tests revealed that, in the inward session, participants significantly increased their variability while the perturbation was applied (+6.0 mels, t = 2.98, p = 0.03, d = 0.77) and maintained a numerically similar increase after perturbation was removed (+4.8 mels, t = 2.28, p = 0.116, d = 0.59). In contrast, participants in the outward session increased their variability in the exposure phase (+3.8 mels, t = 4.84, p < 0.001, d = 1.25), but did not maintain the variability increase when normal feedback was restored (test phase: +0.8 mels, t = 0.58, p = 1.000). As an additional test, we directly compared variability changes across sessions by normalizing the variability to each session’s baseline phase on a participant-specific basis. This analysis showed a reliable main effect of session during both the late exposure (F(2,28) = 6.823, p = 0.004, ηp = 0.328) and test (F(2,28) = 3.422, p = 0.047, η_p_^2^= 0.196) phases. Post-hoc tests confirmed that the baseline-normalized variability significantly differed between inward and control sessions during both the late exposure (t = 2.87, p = 0.037, d = 0.74) and test (t = 2.81, p = 0.042, d = 0.72) phases, while the baseline-normalized variability significantly differed between outward and control sessions during only the late exposure (t = 3.00, p = 0.029, d = 0.77), and not the test phase (t = 0.297, p = 1.000). These results are consistent with those of our previous behavioral study (Tang et al., 2022). Although the variability increase during the late exposure phase in the inward perturbation session was not significantly different from baseline (i.e., 0), unlike the previous study (N=22), it was nonetheless significantly different from the variability change in the same phase (i.e. the late exposure) of the control session.

**Figure 2.**
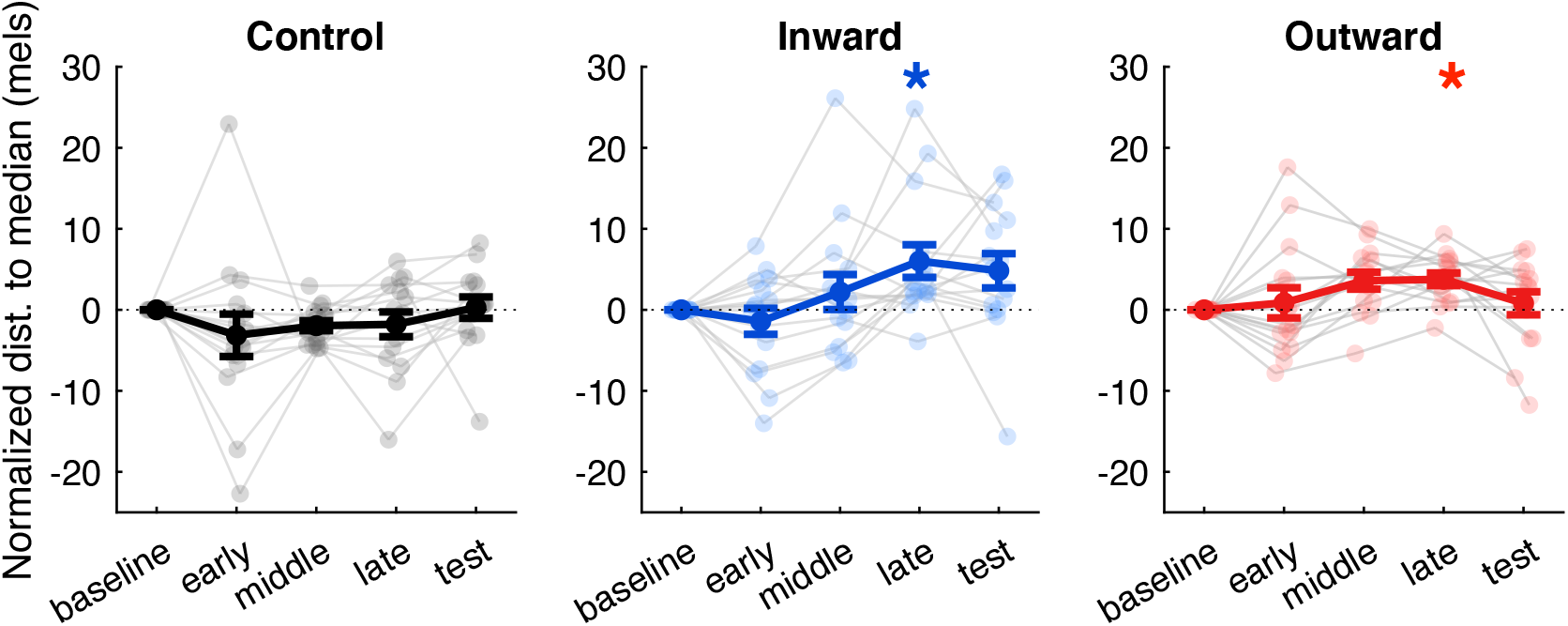
Baseline-normalized variability changes across sessions. (control, inward, and outward). Individual and group means are indicated by thin lines with small transparent dots and thick lines with large solid dots, respectively. Error bars show standard error (SE). * indicates significant change (p < 0.05) from baseline.

In our previous work (Tang et al., 2022), the outward perturbation resulted in an increase not only in produced variability, but also in vowel centering (+2.2 mel), defined as the reduction in variability from vowel onset (first 50 ms) to vowel midpoint (middle 50 ms). Because vowel centering may reflect corrections to ongoing vowel trajectories (Niziolek et al., 2015), this metric may also be related to auditory error sensitivity, although online corrections and trial-to-trial adaptation may be differentially sensitive to errors (Franken et al., 2019; Lester-Smith et al., 2020; Parrell et al., 2017). In the pilot group (N=10), we observed numerically larger (+4.6 mel) but non-significant increases in vowel centering during the outward perturbation session (Figure S1, late exposure phase: t = 1.94, p = 0.251). In the main MEG study, no significant change in centering was observed in participants in any of the three perturbation sessions (outward: -0.41 mel, p > 0.05 in all cases; see supplementary materials).

No participants reported awareness of the perturbation, and none correctly identified the perturbation as a change to their vowels when informed after the final session that their speech had been manipulated (see Table S1).

In sum, we replicated our previous behavioral results: participants exposed to both inward-pushing and outward-pushing perturbations unconsciously increased their produced variability, but maintained this increase when normal auditory feedback was restored only in the inward-pushing condition.

### Alteration of perceived variability affects neural sensitivity to auditory errors

Next, we examined whether the nervous system’s sensitivity to errors, as measured by the magnitude of SIS (see Figure 3 and Methods), can be modulated by auditory perturbations that alter speakers’ perceived variability. Because the magnitude of SIS is modulated by the perceived error (Behroozmand and Larson, 2011, Sitek et al. 2013; Chang et al., 2013; Niziolek, Nagarajan, & Houde, 2013), in the current experiment, increased error sensitivity will be reflected by greater prediction error and, in turn, a decrease in the magnitude of SIS in the test phase compared to the baseline phase.

**Figure 3.**
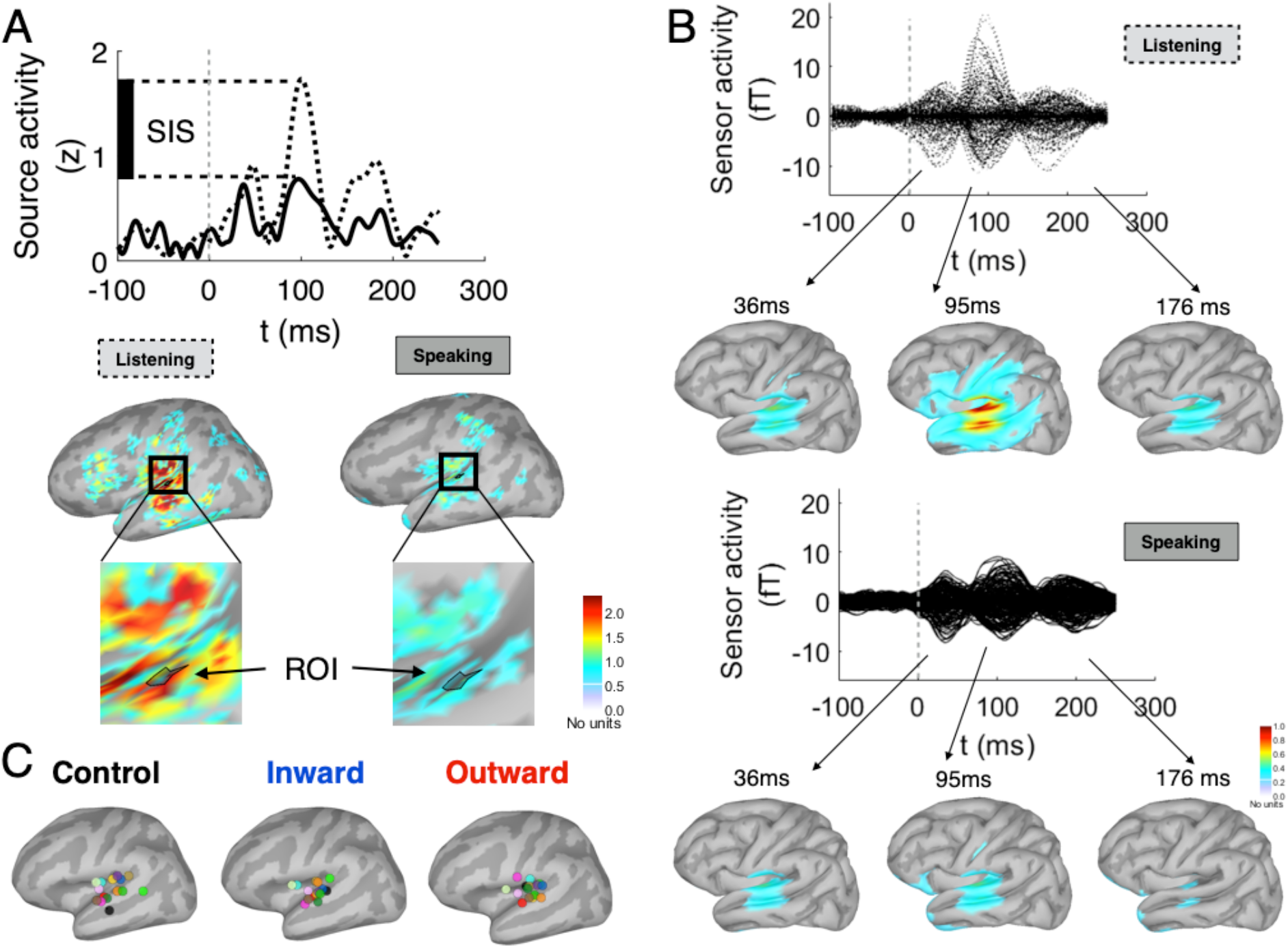
Speaking-induced suppression (SIS) in left auditory cortex. **A**. Upper panel: Time courses of source activity extracted from regions of interest (ROIs, see lower panel) for speaking (solid line) and listening (dashed line) during the baseline phase in a representative participant. SIS is calculated as the difference in M100 amplitude between the listening and speaking peaks (vertical bar on the y-axis). Lower panel: Source cortical maps (dSPM) during listening and speaking trials in the baseline phase at M100 peak (102 ms after sound onset). ROIs were defined as the 10 vertices surrounding the vertex with the largest response (restricted to left temporal and inferior parietal areas) at the time of the M100 peak during listen trials. **B**. Time courses of averaged sensor activity for baseline listening (upper panel, dashed lines) and speaking (lower panel, solid lines) during the baseline phase (averaged across all participants and all three MEG session baselines). Averaged, spatially-smoothed cortical source maps at three visible peaks (36 ms, 95 ms, and 176 ms after sound onset) are shown below. **C**. ROI locations for each of the three MEG sessions projected on a common cortical surface template. Individual participants are indicated by different colors.

All participants exhibited SIS: the M100 peak was suppressed in the speaking task relative to the listening task during all three sessions (Figure 4A). We confirmed that the SIS magnitudes during baseline phase did not differ across sessions (F(2,28) = 0.613, p = 0.549; Figure 4B), suggesting that this measure is stable over time in the absence of any auditory perturbation.

**Figure 4.**
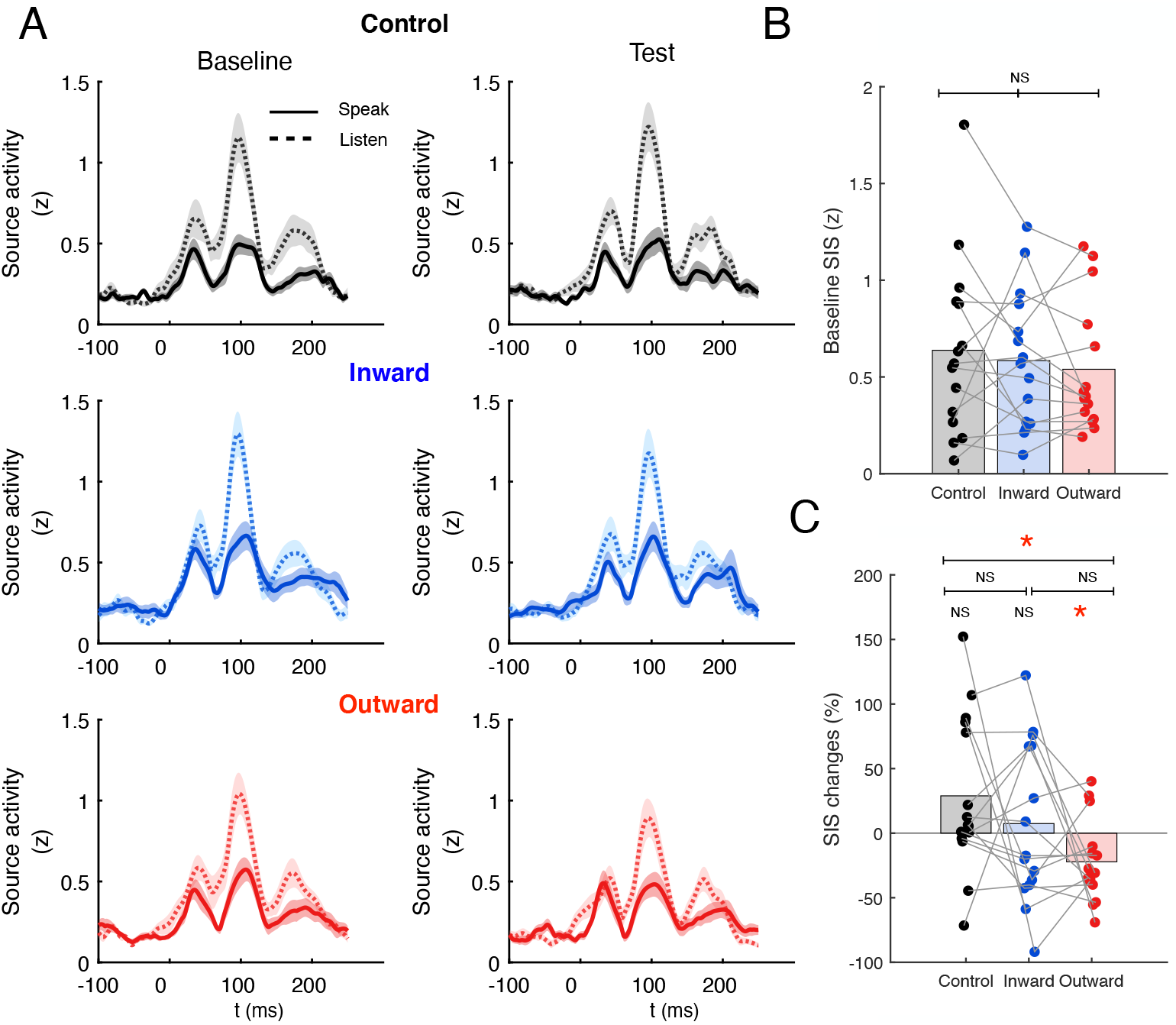
Modulation of SIS across sessions. (control, inward and outward). **A**. Source-localized auditory cortical time course aligned to sound onset during speaking (solid line) and listening (dotted line), shown separately before (baseline, left column) and after (test, right column) exposure to auditory perturbations. Shaded regions around the MEG traces indicate SEM across participants. **B**. SIS magnitudes across sessions obtained during baseline recording. **C**. Normalized SIS changes (%, calculated as 100*(SIS_base_ − SIS_test_)/SIS_base_) across sessions. Group means are indicated by transparent colored bars. Connected points represent data from individual subjects. * indicates significance (p < 0.05).

Consistent with our predictions, SIS was attenuated after exposure to the outward perturbation (decrease of 22.2% ± 8.2 s.e., t = 2.70, p = 0.017, d = 0.69; Figure 4C), suggesting that speakers became more sensitive to auditory errors in this condition. This was true even though produced variability had returned to baseline levels; in other words, the decrease in SIS was not due to greater acoustic error during the test phase of this session. Conversely, SIS increased by 7.4% (± 16.1 s.e., t = 0.46, p = 0.652) after exposure to the inward perturbation, and by 28.9% (± 15.6 s.e., t = 1.85, p = 0.089) in the control session with no perturbation. Normalized SIS changes in the test phase differed significantly across sessions (F(2,28) = 3.237, p = 0.054, η_p_^2^= 0.188). Post-hoc tests confirmed that normalized SIS changes differed between the outward perturbation and control sessions (t = 2.95, p = 0.032, d = 0.76), but not between the inward perturbation and control sessions (t = 0.89, p = 1), nor between the inward and outward perturbation sessions (t = 1.6, p = 0.393).

Although the current study was designed and powered primarily to assess group-level changes in neural markers of auditory error sensitivity, we additionally performed Spearman’s correlation analyses between changes in produced variability and SIS, separately for each session. The results did not show any significant correlation between behavioral and SIS measures (ρ < 0.2, p > 0.05 in all cases). However, this negative result should be interpreted with caution given that: 1) these two measures were necessarily calculated from different blocks in the experiment (i.e., variability was compared between baseline and late exposure phases, while SIS was compared between baseline and test phases; 2) even at 80% power (β=0.2, α=0.0), a sample size of 47 would be required to detect a moderate correlation, suggesting the sample size in the current study (N=15) did not have enough power to reliably detect any potential correlation between our behavioral and neural measures.

### Suppression decreases with acoustic deviance

A previous study found that SIS was reduced in less prototypical productions (i.e., trials farther from the center of the vowel distribution) compared to more prototypical productions (i.e., trials closer to the center of the vowel distribution), suggesting that the auditory system is sensitive to sensory errors resulting solely from motor variability (Niziolek, Nagarajan, & Houde, 2013). Although not the main focus of the current study, we conducted an analysis to confirm these findings in the current dataset. Following previous methods, we divided baseline trials into tertiles based on their distance to the median formants for that vowel: center trials were defined as the closest tertile and peripheral trials as the farthest tertile (see Figure 5B and Methods). Confirming previous work, we found that SIS was smaller in peripheral trials relative to center trials in all three sessions (Figure 5A, C: F(1,14) = 13.179, p = 0.003, η_p_^2^= 0.485). This reduction in SIS for peripheral trials did not differ between sessions (main effect of session: F(2,28) = 2.692, p = 0.085; interaction between trial type and session: F(2,28) = 0.672, p = 0.519).

**Figure 5.**
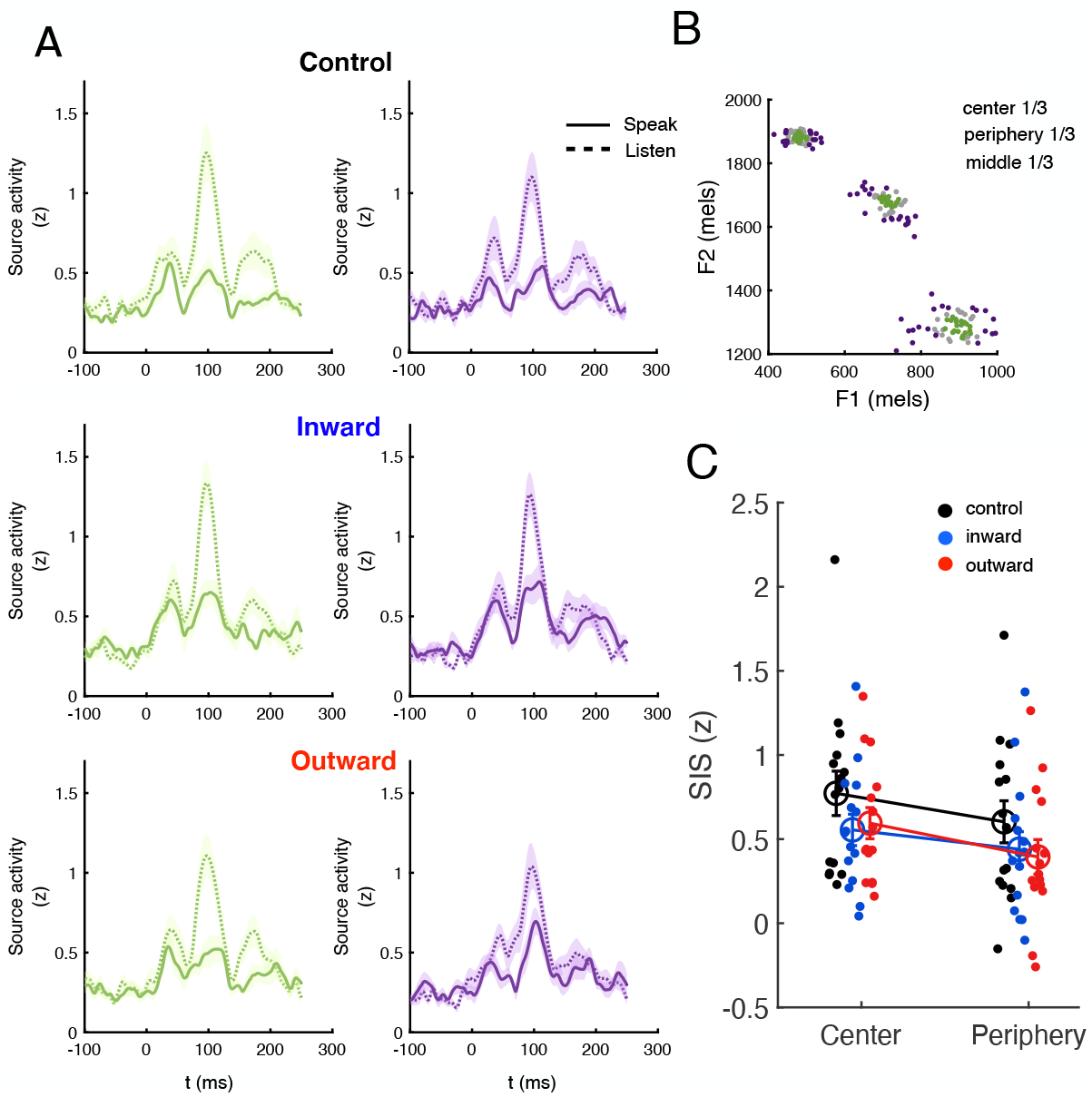
Center vs. periphery vowel productions. **A**. Source-localized auditory cortical time-course aligned to vowel onset, during speaking (solid line) and listening (dotted line), separated into trials at the center (left column) and periphery (right column) of each vowel’s distribution. Shaded regions represent SEM across participants. **B**. Productions from a single participant in the current study, shown in 2D formant frequency space. **C**. SIS amplitudes measured over trials from the center and periphery across sessions (control, inward and outward). Large open circles show group means. Error bars indicate SE. The individual SIS amplitudes are shown in small filled circles.

## Discussion

This study used a widely observed phenomenon in the auditory-motor domain, speaking-induced suppression (SIS), as a window into sensory error processing at the cortical level. Results showed that neural sensitivity to sensory errors, as measured by this suppression, can be modulated by altering speakers’ perceived variability. In particular, error sensitivity was increased after exposure to an outward-pushing perturbation that increased participants’ perceived variability, in line with predictions generated from previous behavioral data and state-space modeling.

Given its importance to sensorimotor learning, sensitivity to sensory error has been extensively studied in the past two decades. Growing behavioral evidence has shown that the sensitivity to sensory error is affected by a number of factors. Individuals adjust their error sensitivity based on the level of confidence they have in their sensory feedback (Burge et al., 2008; Korenberg and Ghahramani 2002). For example, when the visual feedback from a movement outcome is blurry, individuals are less likely to modify their motor commands than when it is sharp (Izawa & Shadmehr, 2008; Wei & Körding, 2010). Avraham et al. (2020) found that error sensitivity increased in consistent environments and suggested that such an increase was mainly due to the contribution of explicit strategies rather than the implicit process driven by sensory prediction errors. Reward and punishment feedback have also been shown to affect the sensitivity to error. Providing explicit rewards can change the speed of adaptation and also enhance the retention of adaptation (Galea et al., 2015; Mawase et al., 2017; Nikooyan & Ahmed, 2015). Again, the impact of reward on sensorimotor adaptation is thought to be mainly exerted by engaging the explicit, cognitive component of adaptation instead of the implicit, unconscious component (Codol et al., 2018). By modulating the implicit cost of error during sensorimotor adaptation, a recent study found that error sensitivity was elevated when the cost of error was large (Sedaghat-Nejad & Shadmehr, 2021). Such implicit error cost affected error sensitivity only and had no significant effect on retention. The novel contribution of our study to the literature on error sensitivity is the demonstration that sensitivity to sensory error can be modulated by alteration of perceived variability during speech adaptation. Compared to visuomotor adaptation during reaching, auditory-motor adaptation is thought to be a more implicit process, occurring without learner awareness (Keough et al., 2013; Kim & Max, 2020; Lametti et al., 2020; Munhall et al., 2009). No participants reported any conscious awareness of the auditory perturbations we applied in the current study or previous behavioural study (Tang et al., 2022). Thus, our results suggest that manipulation of perceived motor variability can modulate sensory error sensitivity during implicit sensorimotor learning.

While evidence from behavioral studies suggests error sensitivity is malleable, there is limited neural evidence regarding whether and how error sensitivity can be modulated in the human brain. In the current study, we used SIS to measure the change in error sensitivity at the cortical level before and after exposure to auditory perturbations that alter speakers’ perceived variability. Exposure to an outward perturbation resulted in an attenuation of SIS during the test phase, after the perturbation had been removed. Importantly, participants’ produced variability had returned to baseline levels during this test phase, suggesting the decrease in SIS can be attributed to an increase in sensitivity to the same sensory error. In the inward perturbation session, we observed no increase in SIS. However, participants’ produced variability remained at elevated levels during the test phase compared with baseline. One possibility, therefore, is that error sensitivity may have decreased as a result of the inward perturbation, but that this decrease was cancelled out by the increase in sensory error they experienced during the test phase. Nevertheless, it is also possible that error sensitivity cannot be modulated by inward perturbation. Future analyses comparing SIS in the trials with the same sensory error (i.e., produced variability) during the inward perturbation session would help clarify this point.

It is important to note that, in this study, SIS was used to measure error sensitivity, but we do not argue that sensory prediction error arises exclusively in the sensory cortex. The cerebellum, an integral part of the motor system, is widely thought to generate predictions and compute and process sensory prediction errors (Kawato, 1999; Diedrichsen et al., 2005; Shadmehr et al., 2010; Tseng et al., 2007). For example, increased cerebellar activation has been found in the presence of prediction errors arising from an unexpected sensory event or the absence of an anticipated somatosensory stimulus (Schlerf et al., 2012). Numerous animal studies have indicated that error signals are encoded in the complex spikes of Purkinje cells of the cerebellum (e.g. Kitazawa et al., 1998; Kobayashi et al., 1998; Streng et al., 2018).

Such prediction errors then modulate a range of neuronal responses (den Ouden et al., 2012). For example, in the auditory-motor domain, the magnitude of SIS decreases in response to an increase in perceived error induced by external auditory perturbation (Chang et al., 2013; Behroozmand and Larson, 2011) or arising from internal variability (i.e., acoustic deviance; Niziolek, Nagarajan, & Houde, 2013). Therefore, although it is not feasible to measure cerebellar-mediated prediction error by directly recording cerebellar output using MEG, we can hypothesize changes in prediction error by assessing SIS at the auditory cortex. It should be noted that SIS, measured in this study using MEG, reflected an average response from left temporal and inferior parietal areas (see Figure 3 for individual ROIs). A previous study using direct cortical recording (ECoG/iEEG), with higher spatial resolution, suggested that SIS and speech perturbation-response enhancement (SPRE) might be encoded at distinct temporal and inferior parietal regions (Chang et al., 2013). This is consistent with single-unit animal data showing that distinct subgroups of sensory neurons are responsible for suppression and enhancement (Eliades & Wang, 2003; Nelson et al., 2013; Schneider et al., 2014).

Several issues remain to be addressed in future work. First, we did not find a significant correlation between behavioral and neural measures (i.e., changes in acoustic variability associated with changes in SIS magnitude). As mentioned in our results, the behavioural and SIS changes were not measured in the same phase. Moreover, the sample size in the current study (N=15) had limited power to detect any potential correlation. It would be important in the future to more directly test the correlation between behavioral and neural measures with a larger sample size. Second, in the main MEG study, in contrast to our previous behavioural study (Tang et al., 2022), we did not observe an increase in vowel centering during the outward perturbation session. In the pilot group, we did observe an increase in centering which was numerically similar to the results of Tang et al. (2022), but the increase was not statistically significant. To observe a reliable centering change, a larger sample size (e.g., >20) seems to be required, which also suggests that centering might be a less sensitive and noisier measure of error sensitivity. Future research is needed to determine the validity and reliability of centering as an index of online error correction.

In summary, error sensitivity, which determines how much we learn from erroneous movements, is a critical factor in sensorimotor learning. Previous work has provided behavioral evidence that error sensitivity can be modulated, but neural evidence has so far been lacking. Here, we took advantage a well-established neural response during speech production that reflects changes in auditory error sensitivity. For the first time, we found the nervous system’s sensitivity to errors can be modulated by altering perceived variability. Enhancing our ability to learn from erroneous movements is a crucial element in practical settings such as rehabilitation.

## Methods

### Participants

Twenty-six native speakers of American English participated: 16 in the main MEG study and 10 in a small behavioral pilot before the main study. The pilot group of 10 participants (7/3 females/males, 18-39 years, mean 23.8 ± 5.9) were recruited to confirm the effect of auditory perturbation on produced variability under the current experimental procedure, including both speaking and active listening trials. All participants were right-handed with no reported history of neurological disorders. Participants’ hearing thresholds were measured using the modified Hughson-Westlake audiogram procedure (Hughson and Westlake, 1944); all participants had normal hearing as defined by thresholds of 25 dB HL or less for frequencies between 250-4000 Hz. All participants gave their informed consent, and the protocol was approved by the Institutional Review Board of the University of Wisconsin–Madison.

Our main analysis compared the magnitudes of SIS before and after auditory feedback perturbation. One participant who did not exhibit suppression during the baseline phase across all three MEG visits was excluded, as we could not assess our main hypothesis in this participant. In total, data from 15 participants (9/6 females/males, 24-65 years, mean 39.3 ± 12.1) were further analyzed.

### Experimental procedure

Each participant completed three separate sessions where they were exposed to different auditory feedback perturbations (either inward-pushing or outward-pushing) or no perturbation at all (control). The order was counterbalanced across participants. There was at least a one-week interval between sessions (mean interval: 10.1 days) to minimize potential carry-over effects in speech from the auditory perturbation.

During MEG recording, participants were seated upright in a sound-attenuated, magnetically shielded recording room and required to complete two types of tasks: a speaking task and a listening task. Stimulus words (“ease”, “add” and “odd”) were pseudorandomly selected and presented on a Panasonic DLP projector (PT-D7700U-K) for 1500 ms, one at a time. The interstimulus interval was randomly jittered between 0.4-1.15 s. These stimulus words contain the same three vowels as in our previous behavioral study (Tang et al., 2022), but start without an onset consonant to avoid movement artifacts in MEG. During the speaking task, participants were instructed to produce the stimulus words while receiving auditory feedback of their own voices through insert earphones. Participants were instructed to keep still and minimize jaw movement during speech, although not at the expense of vowel quality. During the listening task, participants passively listened to the audio recorded in the previous speaking task. They were instructed to keep their eyes open and look at the screen during both tasks.

Each MEG session was divided into three phases (see Figure 1B):

∘ Baseline phase: 60 productions of each word with unaltered auditory feedback, divided into two blocks (30 productions of each word in each block). Each block of the speaking task was followed by a block of the listening task.
∘ Exposure phase: 150 productions of each word with (inward-pushing or outward-pushing) or without (control) perturbation applied, divided into five blocks (30 productions of each word in each block). See real-time auditory perturbation below for more detailed perturbation information. Only the first block of the speaking task in this phase was followed by a listening task.
∘ Test phase: Identical to baseline phase.

After every 45 trials, participants were given a short break (less than 30 s); a long break (1∼2 min) was given every 90 trials. After completing all three MEG sessions, participants were given a brief questionnaire to assess their awareness of the perturbations. The pilot group completed a single session which followed the same experimental procedure as the outward-pushing session. This group received the same instructions (i.e. keep still and minimize jaw movement during speech) as participants did in the main study, though they were not seated in the MEG recording room.

### Real-time auditory perturbation

During the exposure phase, participants were exposed to no perturbation (control) or auditory perturbations that increase (outward-pushing) or decrease (inward-pushing) their perceived variability (Tang et al., 2022). The inward-pushing perturbation shifted every production towards the center of that participant’s distribution for each vowel (i.e. the median F1/F2 values, the vowel “targets”, Figure 1A). The outward-pushing perturbation shifted every production away from these targets. In speech, vowel sounds are defined by resonances in the vocal tract, known as formants. The first two formant frequencies, F1 and F2, are mostly determined by the height and front-back position of the tongue body, respectively, and sufficient to disambiguate different vowel sounds (Ladefoged, 2001). The perturbation magnitude was 50% of the distance, in F1/F2 space, between the current formant values and the vowel targets. Participants’ median F1/F2 values for each vowel were calculated during the baseline phase and subsequently used to calculate the participant-specific perturbation field.

### Apparatus

We used a modified version of Audapter (Cai et al., 2008; Tourville et al., 2013) to record participants’ speech, alter the speech signal when necessary, and play the (potentially altered) signal back to participants in near real time (an unnoticeable delay of ∼18 ms, as measured on our system following Kim & Max, 2020). Speech was recorded at 16 kHz via a lavalier microphone (Shure SM93, modified for compatibility with MEG recording) placed approximately 4-7 cm away from the left corner of mouth. Speech recordings were played back to the participants via MEG-compatible inserted earphone (TIP-300, Nicolet Biomedical, Madison, WI) at a volume of approximately 80 dB SPL. The volume of speech playback varied dynamically with the amplitude of participants’ produced speech.

MEG data were acquired at Froedtert Hospital, Milwaukee, WI, USA, using a 306-channel (204 planar gradiometers and 102 magnetometers) whole-head biomagnetometer system (Vectorview™, Elekta-Neuromag Ltd., Helsinki, Finland). The raw data were acquired with a sampling rate of 2 kHz and high-pass filtered with a 0.03 Hz cutoff frequency. The head position of participants relative to the sensors were determined using four head-position indicators coils attached to the scalp surface, whose locations were digitized using a Polhemus Fastrak system (Polhemus; Colchester, VT), together with three anatomical landmarks (nasion and pre-auricular points) and ∼100 additional scalp points to improve anatomical registration with MRI. Head position was monitored continuously during the entire MEG recording and confirmed for consistency between blocks. Horizontal and vertical eye movements and heartbeats were monitored with concurrent electrooculograms (EOG) and electrocardiogram (ECG) recording.

In a separate MRI session after all three MEG sessions were completed, high-resolution T1- and T2-weighted images of each participant were obtained with a GE Healthcare Discovery MR750 3-T MR system in order to coregister each participant’s MEG activity to a structural image of his or her own brain.

### Acoustic analysis

Acoustic data were pre-processed and analyzed following the procedures previously described in Tang et al., 2022. F1 and F2 of all recorded speech words were tracked offline using wave_viewer (Niziolek & Houde, 2015), an in-house software tool that provides a MATLAB GUI interface to Praat (Boersma & Weenink, 2019). Linear predictive coding (LPC) order and pre-emphasis values were set individually for each participant. Vowel onset and offset were first automatically detected using a participant-specific amplitude threshold. All trials were then checked manually for errors. Errors in vowel onset and offset were corrected by manually labelling these times using the waveform and spectrogram. Errors in formant tracking were corrected by adjusting the pre-emphasis value or LPC order. In total, a limited number of trials (1.2% in Control, 1.4% in Inward-pushing, 1.1% in Outward-pushing) were excluded due to production errors (e.g. if the participant said the wrong word), disfluencies, or unresolvable errors in formant tracks.

The primary goal of the acoustic analysis was to evaluate how variability changed across the different phases of each session. Variability within each experimental phase was measured as the average 2D distance in F1/F2 space from each vowel to the center of the distribution for that vowel in that phase, measured from the first 50 ms of vowel. For offline analysis, the exposure phase was divided into early (the first block of speaking), middle, and late exposure phases (two speaking blocks of each). We also measured vowel centering, a measure of within-trial variability change which is thought to reflect online correction for auditory errors (Niziolek et al., 2013; Niziolek & Kiran, 2018; Niziolek & Parrell, 2021). Centering was calculated as the change in variability from vowel onset (first 50 ms, d_init_) to vowel midpoint (middle 50 ms, d_mid_): C = d_init -_ d_mid_

Previously Niziolek, Nagarajan, & Houde (2013) showed that SIS was reduced when speech was less prototypical (i.e., for trials whose formants were farther from the center of the distribution in 2D formant space for a given vowel), compared to more prototypical speech productions (trials whose formants were closer to this center), Although not the primary focus of the current study, we conducted similar analyses to attempt to replicate these results with our current data. For each participant, speech productions during the baseline phase (60 productions of each stimulus word with unaltered auditory feedback) were divided into center and peripheral trials, defined as the closest and farthest 20 trials from each vowel target (closest and farthest third of trials, see Figure 5C). Combining these across all three stimulus words, 60 center and 60 peripheral trials were defined in the baseline phase of each MEG session for each participant.

### MEG analysis

#### MEG sensor data preprocessing

A temporal variant of signal space separation (tSSS) using MaxFilter software v2.2 (Elekta-Neuromag, Helsinki, Finland) was first performed to remove external magnetic interferences and discard noisy sensors. MEG preprocessing and source estimation were performed with Brainstorm (http://neuroimage.usc.edu/brainstorm/) (Tadel et al., 2011) combined with in-house MATLAB code.

All recordings were manually inspected to detect segments contaminated by large body/head movements or remaining environmental noise sources, which were discarded from further analysis. Heartbeat and eyeblink artifacts were automatically detected from the ECG and EOG traces and removed using signal-space projections (SSP). Projectors were calculated using principal components analysis (PCA: One component per sensor) with Brainstorm default parameters settings (ECG: [-40, +40] ms, [-13-40] Hz; EOG: [-200, +200] ms, [-1.5-15] Hz). In all participants, the principal components (one for heartbeats and one for eyeblinks) that best captured the artifacts’ sensor topography were manually selected and removed. This was sufficient to remove artifact contamination. The preprocessed data (projectors included) were then band-pass filtered between 4 and 40 Hz with an even-order linear phase FIR filter, based on a Kaiser window design (Brainstorm default settings). The 4-Hz high-pass cutoff was applied to filter out low-frequency movement-related artifacts during speech production, improving detection of the M100. Sound onsets were detected offline automatically from the audio channel using an amplitude threshold (i.e., when the amplitude of the signal increased above 1.4 times the standard deviation of the signal over the entire file) and were corrected manually after visual inspection of the waveform. Filtered data were then baseline-corrected using a baseline period from −700 ms to −400 ms relative to sound onset (avoiding potential pre-speech preparatory movement) and segmented into epochs of 1100 ms (−700 to 400 ms relative to sound onset).

#### MEG source estimation

Source reconstruction was performed in Brainstorm using minimum-norm imaging. MEG sensor data were coregistered to individual anatomical MRIs for each participant using common fiducial markers and head shape digitization. For each MEG block, a forward model of neural magnetic fields was computed using the overlapping-sphere method. The positions and orientations of the elementary dipoles were constrained perpendicularly with respect to the cortical surface. The noise covariance matrix, which was used for source estimation, was calculated from a MEG empty room recording (around 3-6 mins.) collected the same day as the participant’s recordings. Dynamic statistical parametric maps (dSPMs), which are a set of z-scores providing spatiotemporal source distribution with millisecond temporal resolution, were estimated by applying Brainstorm’s minimum-norm (MN) imaging approach (see Figure 3A for source activity maps from a representative participant).

#### Measuring SIS

Source maps were averaged across blocks for conditions (task * experimental phase: speak_baseline, listen_baseline, speak_test, listen_test, 180 trials each). For each MEG session of each participant, region-of-interest (ROI) was picked as 10 vertices (constrained) that have the largest response (restricted to left auditory and inferior parietal areas) at M100 peak during listen_baseline. Figure 3C shows the ROIs chosen for each participant during each MEG session projected on a template brain. Source brain activity was extracted from the chosen ROIs and root mean square (RMS) transformed, yielding a time series of positive evoked response. The M100 peak in each condition was defined as the time point of maximal activity between 85 and 120 ms after sound onset; peaks were confirmed by visual inspection. The M100 amplitudes were then calculated as the mean amplitude across a 20-ms window centered at M100 peak for each condition. In alignment with previous studies (Curio et al., 2000; Niziolek, Nagarajan, & Houde, 2013), SIS was calculated by taking the difference in M100 amplitude between the listening and speaking tasks (Figure 2B; SIS_base_ = Listen_base_-Speak_base;_ SIS_test_ = Listen_test_-Speak_test_).

### Statistical analysis

Following similar procedures previously described in Tang et al., (2022), acoustic statistical analyses were performed with repeated-measures ANOVAs and post hoc tests. Data from baseline, late exposure and test phases were included in the repeated ANOVAs separately for the variability and centering results and for each MEG session, with phase and vowel identity as within-subject factors. Baseline-normalized changes (normalized by subtracting the average value in the baseline from the remaining trials) in variability were also compared between sessions (inward, outward and control) using repeated-measures ANOVAs separately for the late exposure and test phases.

To test our main hypothesis (that error sensitivity, measured as SIS magnitude, can be modulated by perturbations that change perceived variability), normalized SIS changes (calculated as [SIS_base_ - SIS_test_]/SIS_base_*100) from all three MEG sessions were included in a repeated-measures ANOVA, with session (inward, outward, control) as a within-subject factor. We also assessed the significance of normalized SIS changes in each session using one-sample t-tests (against value = 0, H_0_ = no change from baseline). Correlations between SIS changes and variability changes were estimated with Spearman’s correlation. To test for potential changes in SIS based on prototypicality (Niziolek, Nagarajan, & Houde, 2013), we compared the SIS between center (SIS_c_ = Listen_c_-Speak_c_) and peripheral trials (SIS_p_ = Listen_p_-Speak_p_) during baseline of all three MEG sessions using repeated-measures ANOVA with distance (center v.s. periphery) and session (inward, outward, control) as within-subject factors.

For all analyses, post-hoc comparisons with Bonferroni correction were conducted in the event of a significant main effect or interaction. The significance level for all statistical tests was set to P ≤ 0.05. Statistical Analyses for both behavioral and MEG data were conducted in R (R Core Team, 2019).

Although it was not the primary focus of the study, analyses of the latency data (M100 peak) were completed using a three-way repeated ANOVA to test for differences in the within-subject variables of task (listening, speaking), session (inward, outward, control), time (baseline, test) and any interactions. See supplementary materials for latency analysis results.

## Supplementary materials

**Figure S1.**
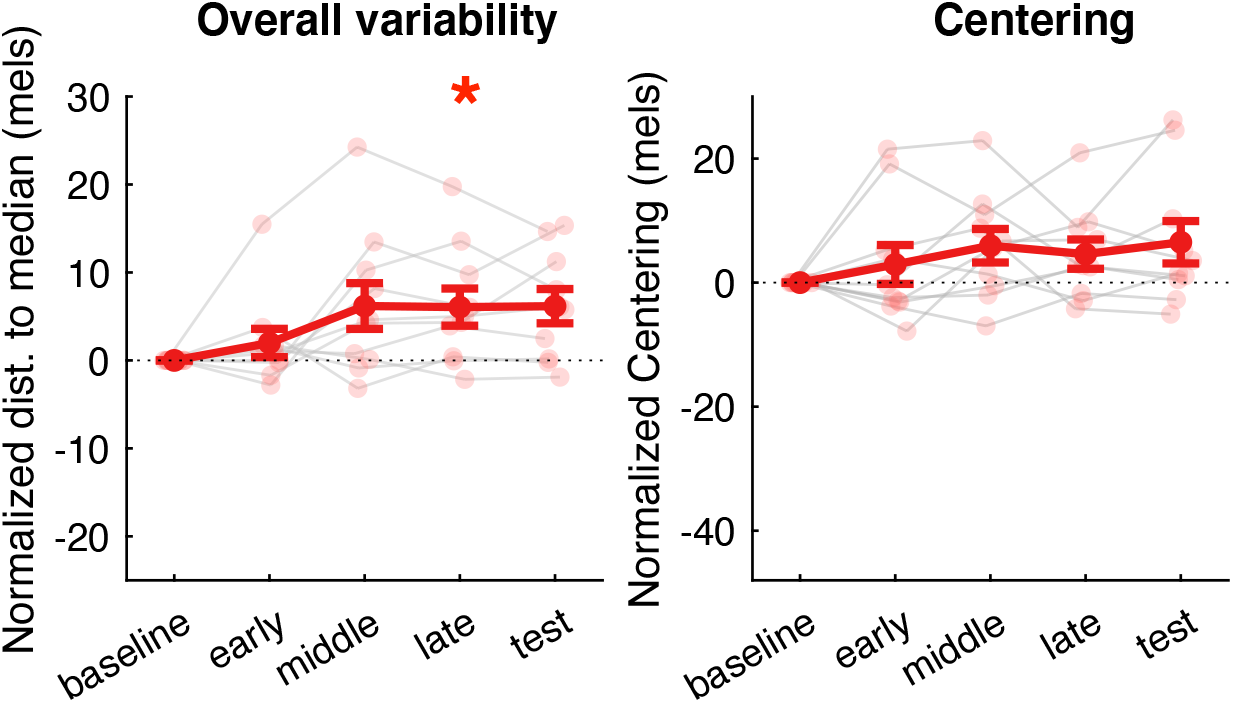
Overall variability (left)and centering (right) changes in pilot behavioural study (normalized by subtracting the average value in the baseline from the remaining trials). Individual and group means are indicated by thin lines with small transparent dots and thick lines with large solid dots, respectively. Error bars show standard error (SE). * indicates significant change (p < 0.05) from baseline.

**Table S1.**
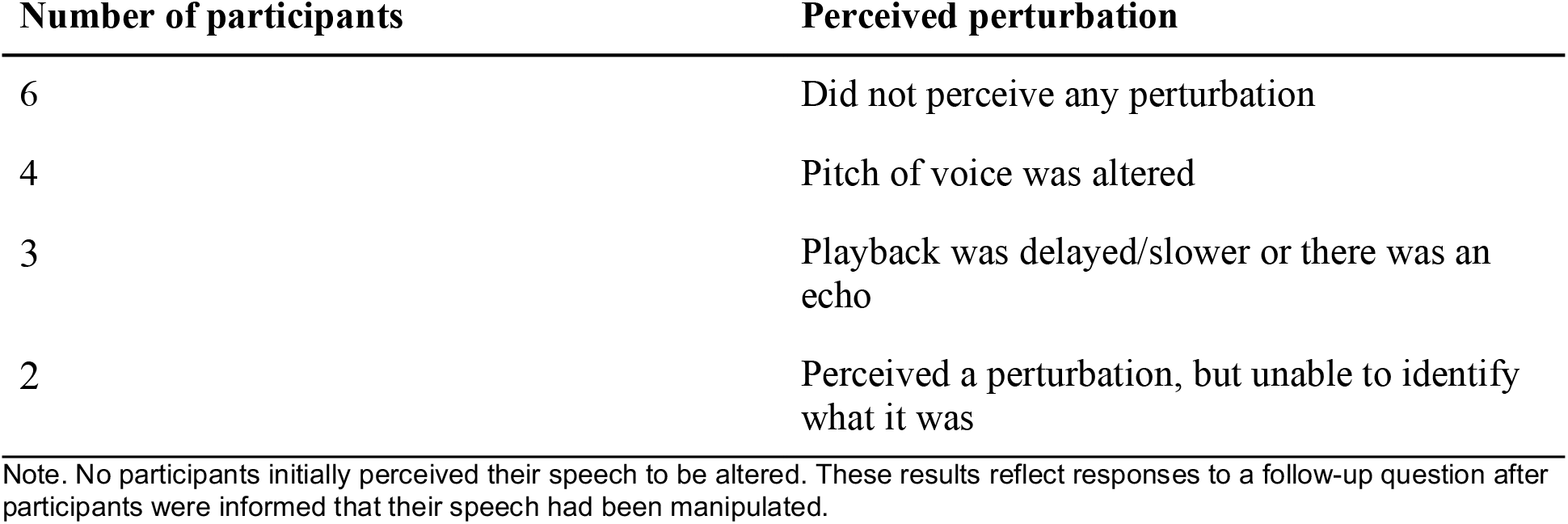
Perturbation awareness

| Number of participants | Perceived perturbation |
| --- | --- |
| 6 | Did not perceive any perturbation |
| 4 | Pitch of voice was altered |
| 3 | Playback was delayed/slower or there was an echo |
| 2 | Perceived a perturbation, but unable to identify what it was |

Note. No participants initially perceived their speech to be altered. These results reflect responses to a follow-up question after participants were informed that their speech had been manipulated.

## Centering results

No significant change in centering was observed in either inward or (Figure S2, main effect of phase: F(2,28) = 0.103, p = 0.903), outward (main effect of phase: F(2,28) = 1.676, p = 0.205), or control (main effect of phase: F(2,28) = 1.239, p = 0.305) session, which was partially inconsistent with our previous finding that an outward perturbation led to an increase in centering during the late exposure phase.

**Figure S2.**
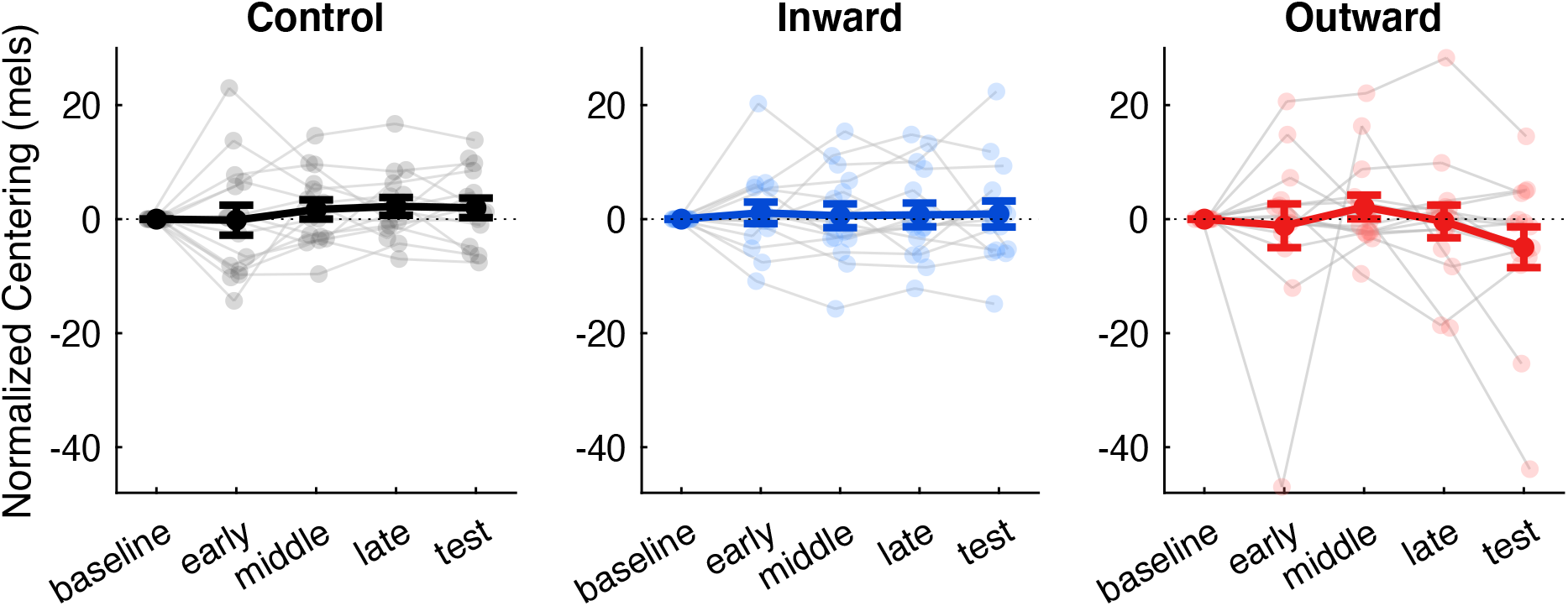
Centering changes (normalized by subtracting the average value in the baseline from the remaining trials) across sessions (control, inward and outward). Individual and group means are indicated by thin lines with small transparent dots and thick lines with large solid dots, respectively. Error bars show standard error.

## M100 latency comparison

A three-way ANOVA with within-subject factors of task (listen, speak), session (in, out, control), time (baseline, test) found a main effect of task (F(1,14) = 7.99, p = 0.013, η_p_^2^= 0.364): M100 latency was slightly but significantly earlier in the listening trials (98 ± 2 ms s.e) than in the speaking trials (102 ± 2 ms s.e), consistent with observations from previous studies (Niziolek, Nagarajan, & Houde, 2013). No differences in M100 latency were found across time (baseline: 101 ± 2 ms s.e, test: 100 ± 2 ms s.e; F(1,14) = 1.77, p = 0.205) or between sessions (in: 99 ± 2 ms s.e, out: 101 ± 2 ms s.e, control: 101 ± 2 ms s.e; F(2,28) = 0.683, p = 0.513), suggesting M100 latency was not affected by any perturbations and kept stable over time. No interactions between these factors were found (p > 0.05 in all cases).

